# CpxR-CpxS two-component system exacerbates development of antibiotic resistance in *Pseudomonas aeruginosa*

**DOI:** 10.64898/2026.10.28.762690

**Authors:** Zhe-Xian Tian, Ai-Ling Tian, Kwan Soo Ko, Yi-Ping Wang

**Affiliations:** State Key Laboratory of Gene Function and Modulation Research, School of Life Sciences, Peking University, Beijing, 100871, China; Southern University of Science and Technology, Shenzhen, Guangdong, 518055, China; Department of Microbiology, Sungkyunkwan University School of Medicine, Suwon, South Korea; Yazhouwan National Laboratory, Sanya, Hainan, 572024, China

**Keywords:** CpxR, CpxS, Two-component system, cross-talk, MexAB-OprM, efflux pump

## Abstract

*Pseudomonas aeruginosa*, as a member of the ESKAPE pathogens, is well known for its ability to develop multidrug resistance following antibiotic therapy. Overexpression of the efflux pumps, often caused by dysfunction of related transcriptional factors, is well known as a common mechanism underpins multidrug resistance. Previous work showed direct regulatory link of CpxR-CpxS, an OmpR-EnvZ family two-component system, with the expression of the predominant MexAB-OprM efflux pump in *P. aeruginosa*. However, the contribution of this regulatory link to the development of antibiotic resistance has not been fully elucidated. In this work, it has been shown that deletion of *cpxR*, but not *cpxS*, reduces the emergence of multidrug resistant colonies almost 1000 times lower compared to the parental wild type *P. aeruginosa* PA14 following sub-lethal antibiotic exposure, suggesting that apart from CpxS, other sensory component(s) or pathway(s) can also mediate CpxR activity in the development of antibiotic resistance in *P. aeruginosa*. We further demonstrate that when glucose is used as the sole carbon source, it leads to increased expression levels of *mexA* and reduced antibiotics susceptibilities in the PA14Δ*cpxS* strain in a *cpxR* dependent manner. GtrS, another EnvZ family sensor kinase, but not the well-known AckA-Pta pathway, mediates this glucose effects. Both *in vivo* and *in vitro* evidences support that GtrS can activate CpxR in a cross-talk manner in the absence of CpxS. Besides *gtrS*, at least four other sensor kinase genes (*envZ*, *tctE*, *phoR* and *colS*) can exert activation effect upon CpxR in the PA14Δ*cpxS* strain. Having the risk of multiple way to antibiotic resistance development, *cpxS* null mutations can be emerged in *P. aeruginosa* clinical isolates. Since many OmpR-EnvZ family two-component system have been shown to be engaged in antibiotic resistance, its potential risk of exacerbating antibiotic resistance in ESKAPE pathogens has been discussed.

## Introduction

Antibiotic resistance emerges as one of the most critical public health threats, invalidating current available treatments against bacterial infections(1, 2). The Infectious Diseases Society of America has highlighted the ESKAPE pathogens (*Enterococcus faecium*, *Staphylococcus aureus*, *Klebsiella pneumoniae*, *Acinetobacter baumannii*, *Pseudomonas aeruginosa* and *Enterobacter* spp.) capable of ‘escaping’ the biocidal action of antibiotics(3). Among them, *P. aeruginosa*, a major pathogen associated with cystic fibrosis, is well known for its intrinsic resistance to a wide range of antimicrobial agents and its ability to develop multidrug resistance following antibiotic therapy(4).

Multidrug resistant phenotypes can arise from the boost of intrinsic efflux activities through the overexpression of the efflux pumps, as a common mechanism in bacteria(5). In *P. aeruginosa*, the MexAB-OprM efflux pump, as the predominant multidrug efflux pump, is largely responsible for intrinsic and acquired multidrug resistance(6). The expression of the *mexAB-oprM* operon was modulated directly by two repressors MexR and NalD and indirectly by another repressor NalC. Mutations causing defective forms of MexR, NalC and NalD lead to overexpression of the *mexAB-oprM* operon and enhanced multidrug resistance in *P. aeruginosa*(7–9). Besides these transcriptional repressors, we have found that CpxR, a response regulator of two-component system, can bind and activate the promoter of the *mexAB-oprM* operon and contributes to a novel type of multidrug resistance in *P. aeruginosa*(10). This regulatory link of a two-component system to the expression of the predominant multidrug efflux pump is unique in *P. aeruginosa* among the *Pseudomonas* species, probably achieved by genomic relocation at the promoter region of *mexAB-oprM* operon(10).

CpxR and its cognate membrane integral sensor kinase CpxS mediates various cell envelop stress signals and influences gene expression related to antibiotic resistance, biofilm matrix production, iron acquisition, and redox homeostasis in *P. aeruginosa*(11). As a member of EnvZ sensor kinase family, CpxS detects envelope stresses by the periplasmic domain and delivers the signal through the HAMP domain and autophosphorylates the conserved histidine residue of the Dhp domain and finally transfers the phosphoryl group to CpxR which, in turn, functions as a global transcription activator for the CpxR regulon, which is a common feature of Cpx systems in many bacteria(12). So far, no physiological cues have been found to activate intact Cpx systems strong enough to overcome the inhibitory functions of the multiple transcriptional repressors and elevates the expression of the *mexAB-oprM* operon in *P. aeruginosa*.

The importance of the regulatory link of CpxR-CpxS to the expression of the *mexAB-oprM* operon in terms of the contribution to multidrug resistance had been shadowed, until we discovered the mutational *cpxS* resistome, namely various single amino-acid residue substituted CpxS variants showing constitutive activity can lead to elevated expression level of the *mexAB-oprM* operon and increased multidrug resistance in *P. aeruginosa*(13). Constitutively active mutation often exists in the EnvZ family sensor kinases(14–16), but it has been the first case associated with antibiotic resistance development in bacteria.

In this work, we further investigate how this regulatory link reshapes the physiologic and genetic landscapes of antibiotic resistance development in *P. aeruginosa*. We conducted antibiotic treatments of sub-lethal concentrations and observed the different contributions of CpxR and CpxS in exacerbating antibiotic resistance in *P. aeruginosa*. During the investigation of alternative pathways mediated by CpxR other than CpxS, we observed the existences of cross-talk effects of multiple EnvZ family sensor kinases upon CpxR, which can lead to elevated expression of the *mexAB-oprM* operon and increased multidrug resistance in *P. aeruginosa*. As cross-talk effects often exist among the EnvZ family sensor kinases, its potential risk of exacerbating antibiotic resistance in ESKAPE pathogens has been discussed in this work.

## Results

### Deletion of *cpxR* reduces development of antibiotic resistance in *P. aeruginosa*

Recently, it has been demonstrated that various stress factors, including oxidative stress, trigger the Cpx response in *P. aeruginosa*(11). Earlier studies identified a common pathway of cell death caused by bactericidal antibiotics, which involves the generation of highly harmful hydroxyl radicals within bacterial cells(17). These hydroxyl radicals can also cause genetic mutations. In fact, it has been shown that exposure to sub-lethal doses of antibiotics induces oxidative stress, leading to increased mutation rates and the emergence of multidrug resistance in bacterial populations(18–20). This led us to explore whether CpxR-CpxS plays a role in the development of antibiotic resistance when *P. aeruginosa* is exposed to sub-lethal concentrations of antibiotics. Wild type PA14, PA14Δ*cpxR* mutant, PA14Δ*cpxS* mutant and PA14Δ*cpxS*Δ*cpxR* double mutant strains were exposed to sub-lethal doses of ofloxacin and carbenicillin respectively in 3-round subcultures. Then, the cells were serially diluted and spotted on agar plates to evaluate the development of antibiotic resistance. The CFUs of PA14Δ*cpxR* grown on plates with lethal concentrations of carbenicillin plus ofloxacin were almost 1000 times lower than those of the wild type PA14 in both cases of ofloxacin and carbenicillin sub-lethal concentration treatments (Fig. 1). These results indicate that CpxR exacerbates the development of multidrug resistance in *P. aeruginosa* populations following sub-lethal antibiotic exposure.

**Fig. 1.**
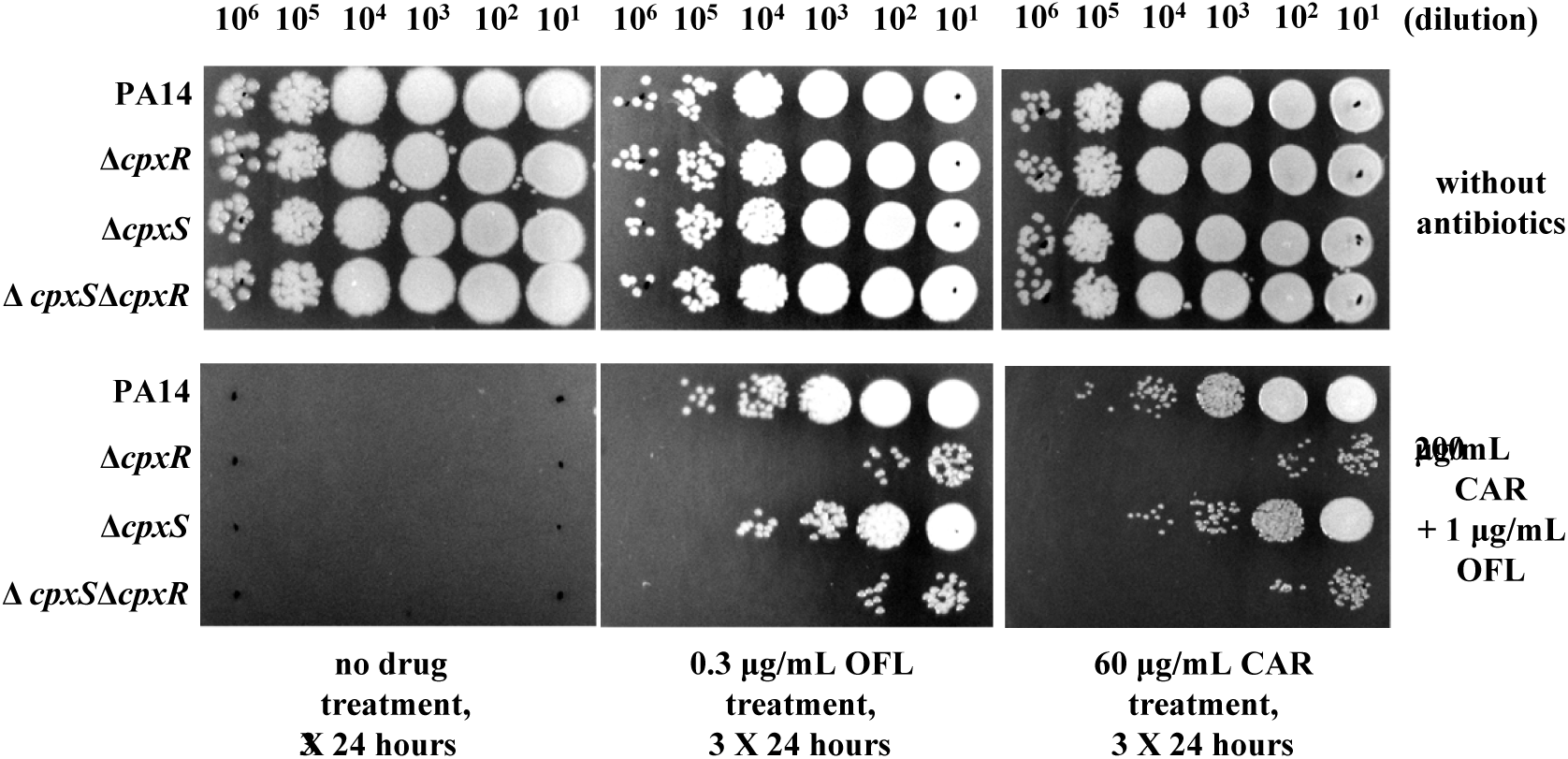
Deletion of *cpxR*, but not *cpxS*, drastically reduces development of multidrug resistance after the 3-round subcultures (3 × 24 hours) exposed to sub-lethal concentration of antibiotics in *P. aeruginosa* PA14. All strains were grown in LB medium at 37 °C with high-speed agitation (250 rpm) and the sub-lethal concentrations of ofloxacin and carbenicillin were 0.3 μg/mL and 60 μg/mL respectively, while the MICs of ofloxacin and carbenicillin were 0.5 μg/mL and 128 μg/mL respectively. After 3-round subcultures, the samples were serially diluted and spotted 2.5 μL on LB agar plates containing lethal concentration of carbenicillin (200 μg/mL) and ofloxacin (1 μg/mL). Each experiment was independently repeated three times and presented one of the similar results.

Surprisingly, when sublethal antibiotic treatments were performed to compare the *cpxS* deletion strain PA14Δ*cpxS* with the wild-type strain PA14, only ten times lower developments of multidrug resistance were observed. Further deleting *cpxR* resulted in drastically decreased development of multidrug resistance (Fig. 1). These results suggest that CpxR may mediate other pathways engaged in the development of antibiotic resistance in the PA14Δ*cpxS* strain.

### Glucose causes reduced antibiotic susceptibility in *cpxS* deleted *P. aeruginosa* strain in a *cpxR* dependent manner

During the identification of CpxR mediated signals, we found that the expression levels of *cpxP* and *mexA* were highly increased in the PA14Δ*cpxS* strain compared to the wild type strain PA14, when the strains were grown in MOPS buffered minimal medium as glucose as the sole carbon source (Fig. 2). Ectopic expression of *cpxS* by introduced plasmid could restore to the expression levels of the wild type strain in the PA14Δ*cpxS* strain. The elevation of the expression levels of *cpxP* and *mexA* was completely abolished, when *cpxR* was deleted in the PA14Δ*cpxS* strain (Fig. 2) Furthermore, the elevation of the expression levels of *cpxP* and *mexA* was also abolished, when glucose was replaced by tryptone in the medium. Accordingly reduced antibiotic susceptibility was observed in the PA14Δ*cpxS* strain grown in the glucose medium compared to the wild type strain (Table 1). Taken together, glucose as the sole carbon source led to increased expression levels of *cpxP* and *mexA* and reduced antibiotics susceptibilities in the PA14Δ*cpxS* strain in a *cpxR* dependent manner.

**Fig. 2.**
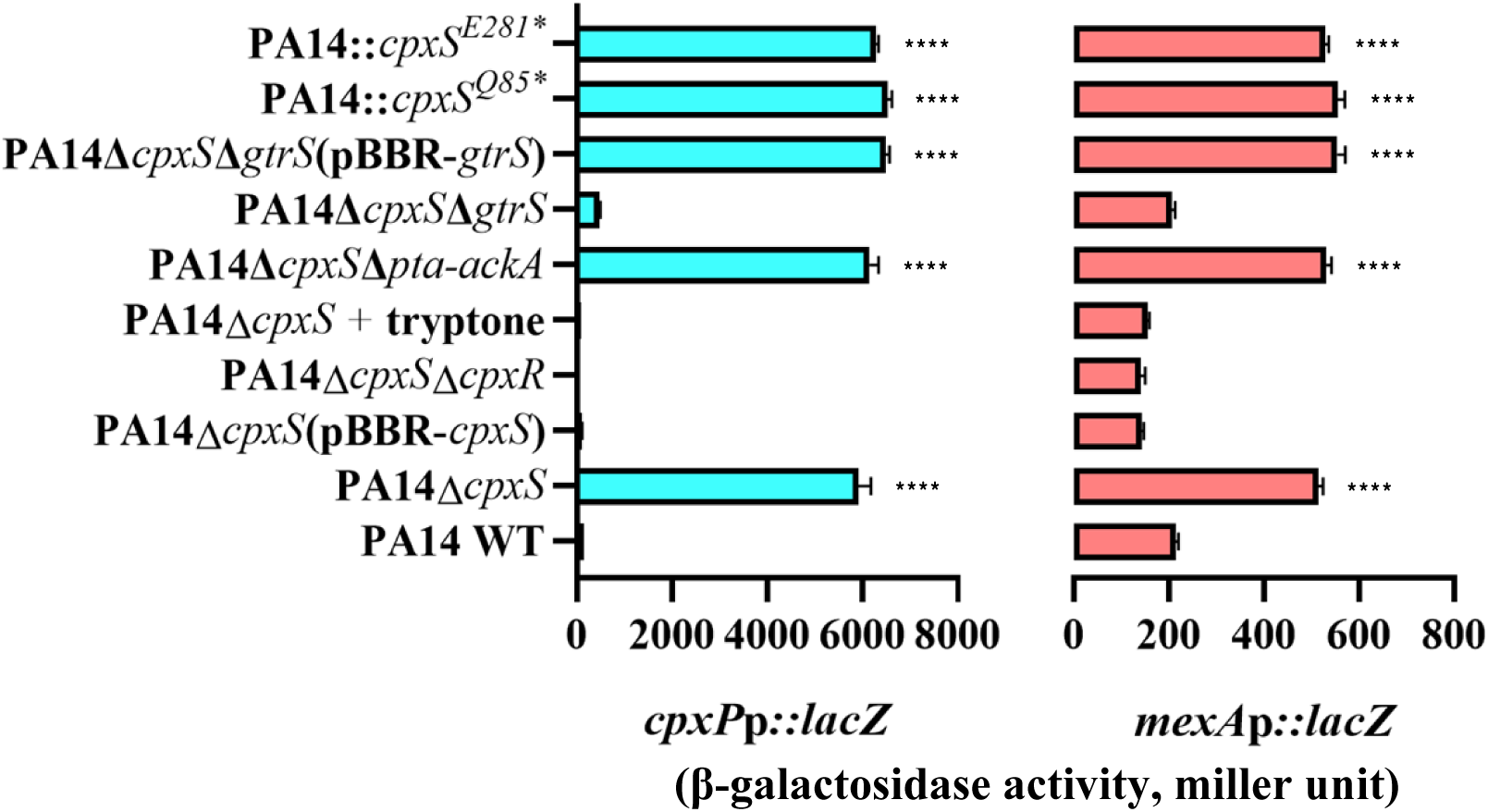
Glucose as sole carbon source causes increased expression levels *cpxP* and *mexA* in *cpxS* null *P. aeruginosa* strains grown in dependence of *cpxR* and *gtrS*, but not *pta-ackA*. Cells containing *cpxP*p-*lacZ* or *mexA*p-*lacZ* reporter plasmids were grown to exponential phase in MOPS buffered minimal medium at 37 °C. β-galactosidase assays were performed and the results were expressed as the mean values from triplicate samples.

**Table 1.**
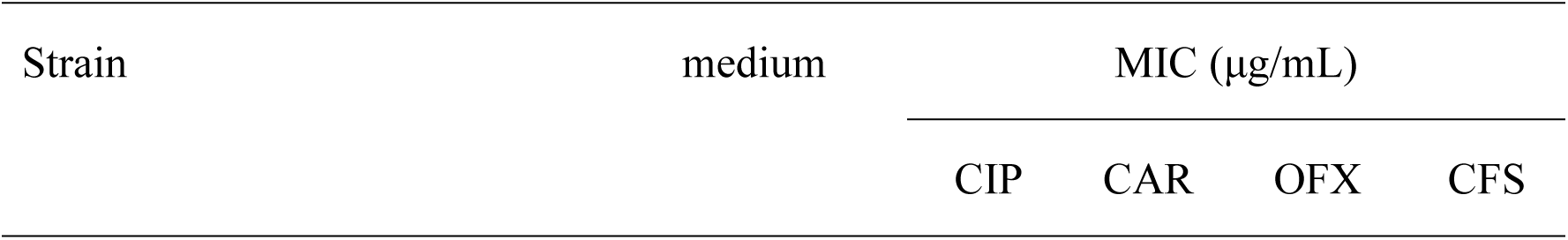
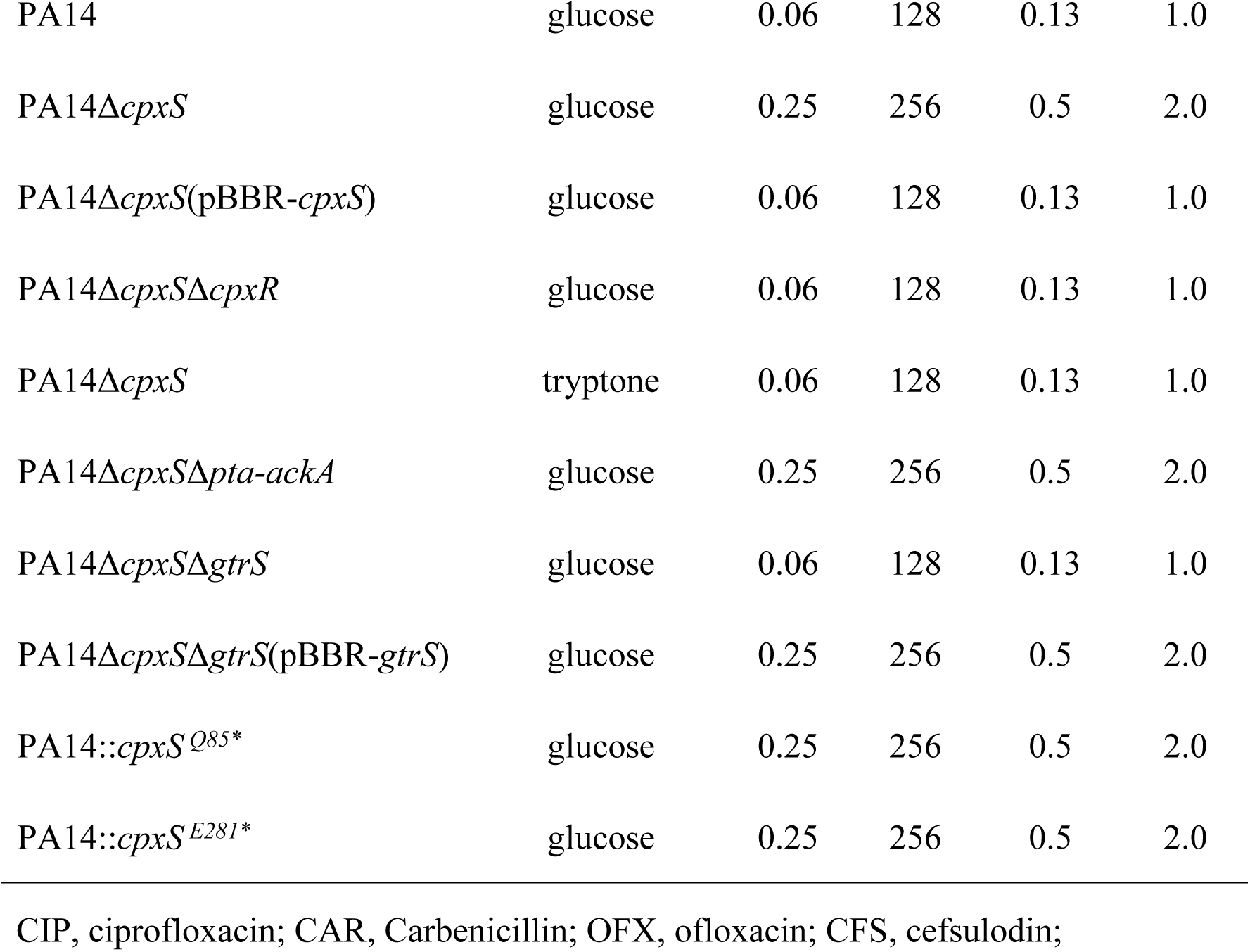
Minimal inhibitory concentrations of antibiotics of *P. aeruginosa* strains grown in different conditions.

| Strain | medium | MIC (μg/mL) |  |  |  |
| --- | --- | --- | --- | --- | --- |
|  |  | CIP | CAR | OFX | CFS |
| PA14 | glucose | 0.06 | 128 | 0.13 | 1.0 |
| PA14Δ*cpxS* | glucose | 0.25 | 256 | 0.5 | 2.0 |
| PA14Δ*cpxS*(pBBR-*cpxS*) | glucose | 0.06 | 128 | 0.13 | 1.0 |
| PA14Δ*cpxS*Δ*cpxR* | glucose | 0.06 | 128 | 0.13 | 1.0 |
| PA14Δ*cpxS* | tryptone | 0.06 | 128 | 0.13 | 1.0 |
| PA14Δ*cpxS*Δ*pta-ackA* | glucose | 0.25 | 256 | 0.5 | 2.0 |
| PA14Δ*cpxS*Δ*gtrS* | glucose | 0.06 | 128 | 0.13 | 1.0 |
| PA14Δ*cpxS*Δ*gtrS*(pBBR-*gtrS*) | glucose | 0.25 | 256 | 0.5 | 2.0 |
| PA14::*cpxS*<sup>Q85*</sup> | glucose | 0.25 | 256 | 0.5 | 2.0 |
| PA14::*cpxS*<sup>E281*</sup> | glucose | 0.25 | 256 | 0.5 | 2.0 |

CIP, ciprofloxacin; CAR, Carbenicillin; OFX, ofloxacin; CFS, cefsulodin;

### Sensor kinase GtrS mediates the reduction of antibiotic susceptibility by glucose in *cpxS* deleted *P. aeruginosa* strain

It has been known that acetyl phosphate, the intermediate of the phosphotransacetylase (Pta)-acetate kinase (AckA) pathway, can activate the transcription of *cpxP* in a CpxA-independent manner by donating its phosphoryl group to CpxR in *E. coli* (21). Recently, it has been reported that Pta-AckA dependent activation of CpxR increased the antibiotic resistance of *cpxA* deletion mutants in *Salmonella enterica serovar Typhimurium* (22). The intracellular level of acetyl phosphate was known to be elevated in *E. coli* grown in glucose(23). In order to investigate whether the elevation of *cpxP* and *mexA* expression levels and the reduction of antibiotic susceptibility by glucose was mediated by acetyl phosphate, the *pta-ackA* loci was deleted in the PA14Δ*cpxS* strain. The *cpxP* and *mexA* expression levels and the antibiotic susceptibility showed no change in the PA14Δ*cpxS*Δ*pta-ackA* strain compared to the PA14Δ*cpxS* strain grown in glucose minimal medium (Fig. 2, Table 1). These results demonstrated that the observed glucose effects were independent of the *pta-ackA* loci and mediated by alternative pathway in the PA14Δ*cpxS* strain.

Previously it was known that GltR-GtrS two-component system was involved in glucose transport and metabolism in *P. aeruginosa*(24–26). In order to investigate if the sensor kinase GtrS mediated the above glucose effect, *gtrS* was deleted in the PA14Δ*cpxS* strain. Then the elevation of *cpxP* and *mexA* expression levels and the reduction of antibiotic susceptibility by glucose were completely abolished in the PA14Δ*cpxS*Δ*gtrS* strain (Fig. 2, Table 1). Ectopic expression of *gtrS* by introduced plasmid could reproduce the increased expression levels of *cpxP* and *mexA* as well as the reduction of antibiotic susceptibility by glucose in the PA14Δ*cpxS*Δ*gtrS* strain (Fig. 2, Table 1). Taken together, sensor kinase GtrS but not AckA-Pta pathway mediates the elevation of *cpxP* and *mexA* expression levels and the reduction of antibiotic susceptibility by glucose in *cpxS* deleted *P. aeruginosa* strain.

GtrS and CpxS belong to EnvZ family sensor kinase and many members of EnvZ family sensor kinase can exert cross-talk effect, namely a sensor kinase activates a non-cognate response regulator in the absence of its sensor kinase partner (27–29). The above observation of GtrS-mediated glucose effect prompted us to investigate if GtrS exert cross-talk effect on the response regulator CpxR in the absence of CpxS in *P. aeruginosa*. When a plasmid harboring gene encoding the cytoplasmic part of GtrS (GtrSc) was introduced, the expression level of *cpxP* was significantly increased in PA14Δ*cpxS* strain grown in medium supplemented with tryptone. This increasement was *cpxR* dependent, as it was abolished by deletion of *cpxR* (Fig. 3A). Since phosphorylated CpxR protein can specifically bind its target DNA(10), it was investigated if the purified GtrSc enable the purified CpxR proteins to bind the target DNA in electrophoretic mobility shift assay. A specific shifted band was observed when GtrSc was co-incubated with CpxR (Fig. 3B). Taken together, both *in vivo* and *in vitro* data demonstrated that GtrS could activate CpxR in a cross-talk manner in the absence of CpxS.

**Fig. 3.**
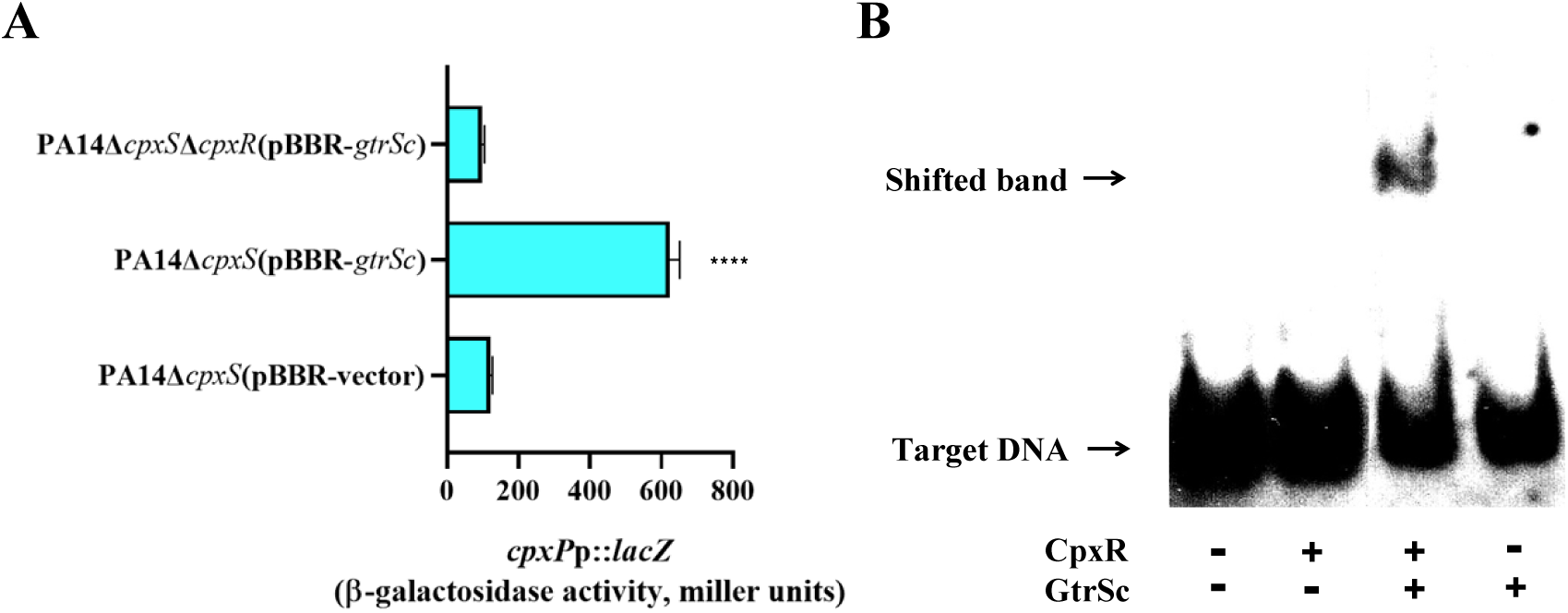
Verification of cross-talk effect between the cytosolic part of GtrS (GtrSc) and CpxR by *in vivo* (A) and *in vitro* (B) experiments. A. The expression level of *cpxP* is increased in the *cpxS* null *P. aeruginosa* strain containing the GtrSc expression plasmid grown in tryptone medium in CpxR dependent manner. β-galactosidase assays were performed and the results were expressed as the mean values from triplicate samples. B. Purified His-tagged CpxR pre-incubated with purified His-tagged GtrSc directly bind to DIG-labelled DNA fragments of *mexA*p *in vitro* illustrated by EMSAs. CpxR proteins were pre-incubated with GtrSc proteins (1:1) in phosphorylation reaction and subsequently CpxR protein (100 nM) and DIG-labelled DNA fragments (0.2 nM) were added to the binding reaction and separated by electrophoresis as CpxR or GtrSc alone and no proteins as controls.

### Multiple sensor kinases can exert activation effect upon CpxR in *cpxS* deleted *P. aeruginosa* strain

*P. aeruginosa* strains have more than 20 OmpR-like response regulator coding genes in its genome (24 in *P. aeruginosa* PAO1 genome), much more than *E. coli* strain (14 in *E. coli* MG1655 genome) (30). Most of them have their cognate EnvZ family sensor kinases, mediating transduction of specific signals, which yet to be fully elucidated. Ectopic expression of *gtrS* by introduced plasmid increased *cpxP* expression level even in the tryptone medium, independent of glucose, in the PA14Δ*cpxS* strain, but not in the PA14Δ*cpxS*Δ*cpxR* strain (Fig. 4), indicating signal bypassed cross-talk by overexpression of *gtrS*. Adopting this advantage of signal bypassing, a series of plasmids harboring EnvZ family sensor kinase coding genes of *P. aeruginosa* PA14 were introduced to investigate if these EnvZ family sensor kinases could exert similar cross-talk effect upon CpxR in the PA14Δ*cpxS* strain. Besides *gtrS*, at least four sensor kinase genes (*envZ*, *tctE*, *phoR* and *colS*) could lead to increased *cpxP* expression level in the PA14Δ*cpxS* strain in the tryptone medium in a CpxR dependent manner (Fig. 4). These data demonstrated that multiple sensor kinases can exert activation effect upon CpxR in the *cpxS* deleted *P. aeruginosa* strain.

**Fig. 4.**
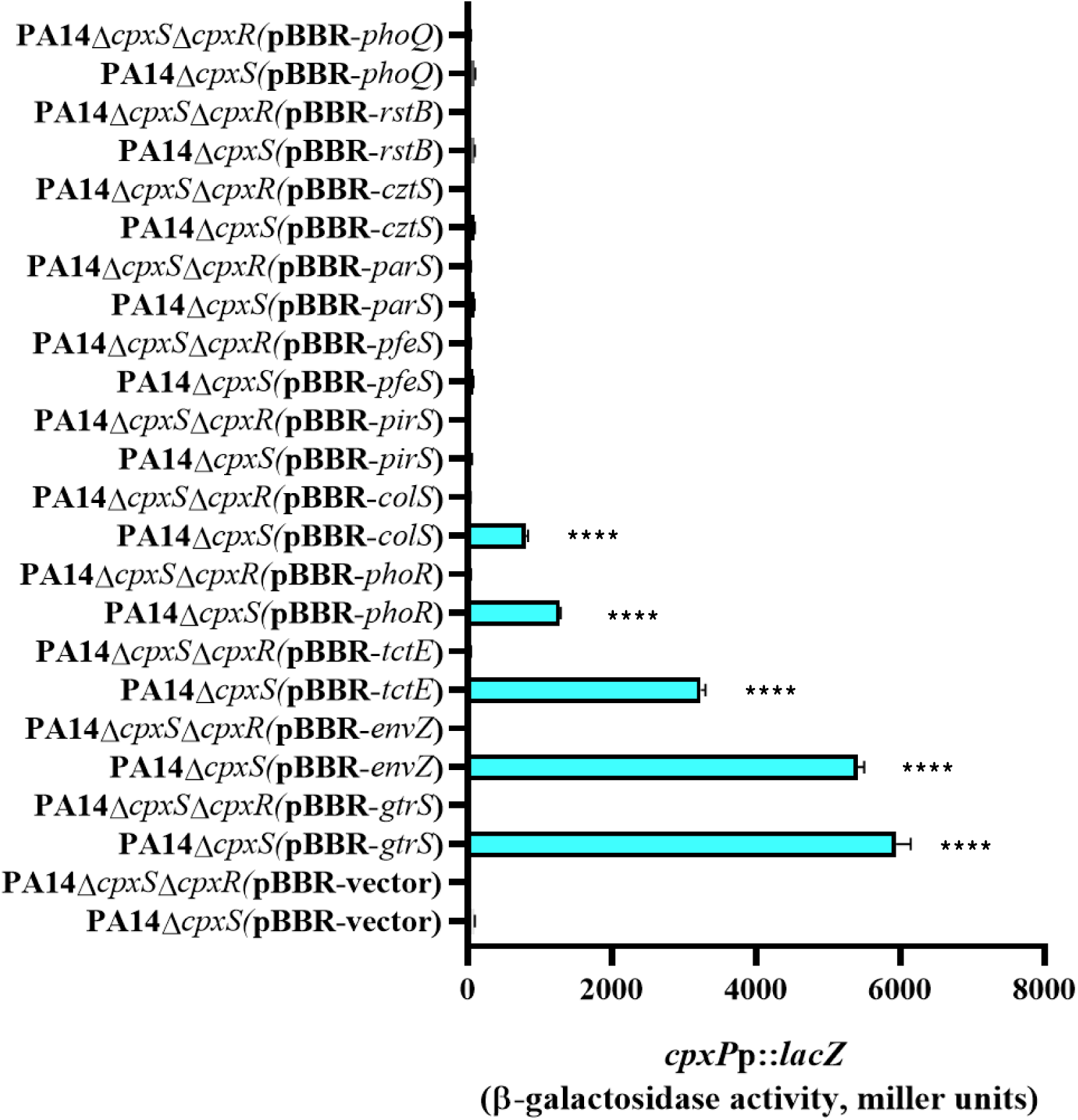
Multiple sensor kinases can exert activation effect upon CpxR in *cpxS* deleted *P. aeruginosa* strain. *In trans* expression of GtrS, EnvZ, TctE, PhoR and ColS by introduced plasmid leads to increased expression level of *cpxP* in *cpxS* deleted *P. aeruginosa* strain. Cells containing *In trans* sensor kinase expression plasmid and *cpxP*p-*lacZ* reporter plasmid were grown to exponential phase in tryptone medium at 37 °C. β-galactosidase assays were performed and the results were expressed as the mean values from triplicate samples.

### Existence of *cpxS* null mutations in *P. aeruginosa* genomes

As *cpxS* null mutation raised a risk of multiple way to the development of antibiotic resistance, it was worth to investigate existence of *cpxS* null mutations in *P. aeruginosa* genomes, especially in clinical isolates. In order to investigate the presence of relevant *cpxS* mutations in *P. aeruginosa* genomes, we searched around 4000 published genomes of *P. aeruginosa* isolates in the Pseudomonas Genome Database(31). Setting the genome of *P. aeruginosa* PA14 strain as reference, the BLASTN analysis revealed that *cpxS* variants strikingly outnumbered *cpxR* variants among these *P. aeruginosa* genomes (Fig. 5, S1 Table). It seems that many sensor kinase variants outnumber their cognate response regulator variants among these *P. aeruginosa* genomes (S1 Fig., S1 Table).

**Fig. 5.**
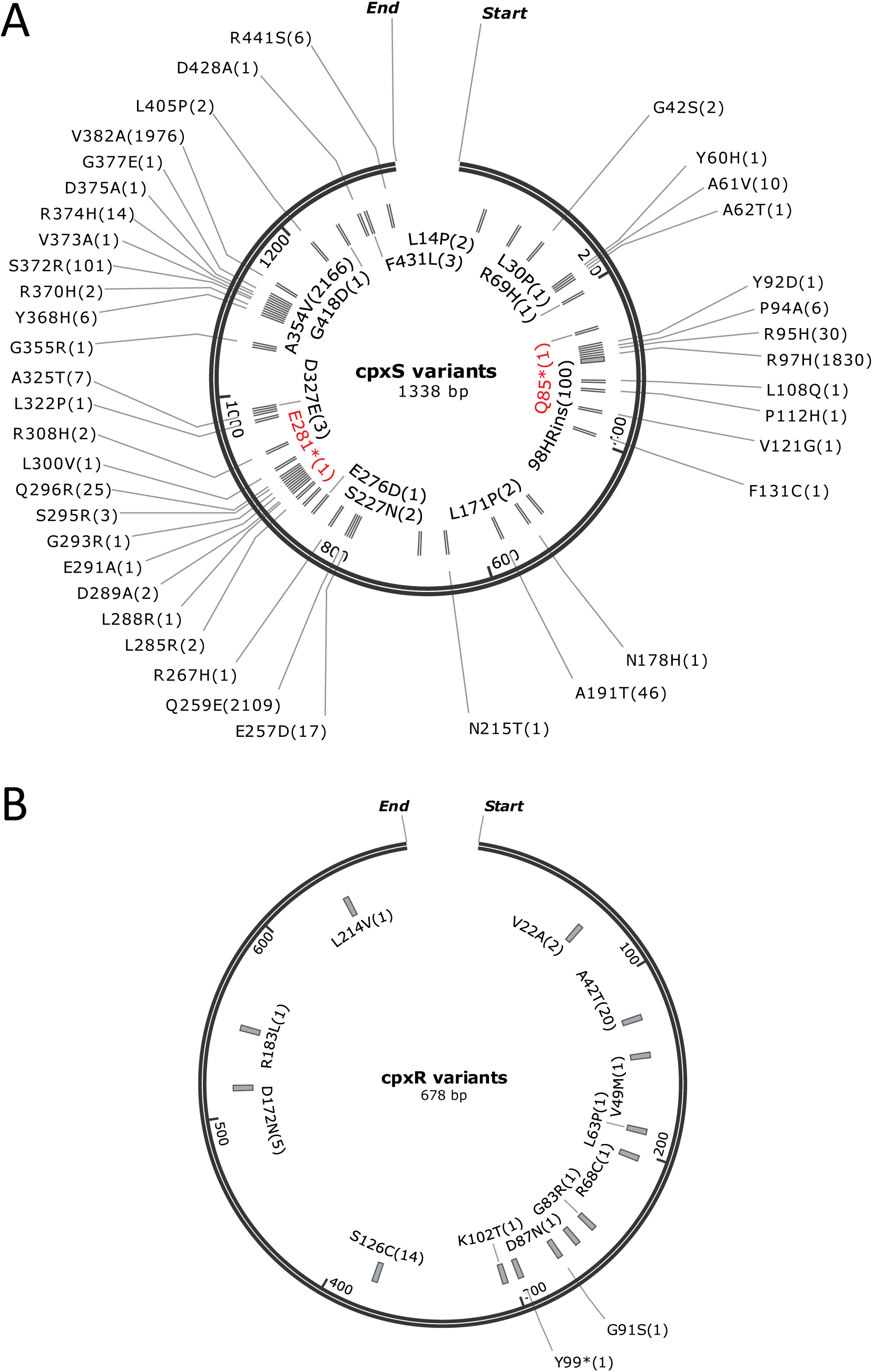
The amino acid substitutions of CpxS variants (A) and CpxR variants (B) in the published genomes of *P. aeruginosa* isolates (S1 Table). Numbers in the parentheses are the occurrence times in different genomes. Nonsense CpxS variants are marked in red.

Among the *cpxS* variants, two nonsense mutations (Q85*, E281*) occurred in an extensively drug-resistant (XDR) *P. aeruginosa* clinical isolate(32) (Accession No. NZ_CAADLG010000522.1 in S1 Table) and a β-lactam/β-lactamase inhibitor resistant *P. aeruginosa* clinical isolate(33) (Accession No. NZ_JADSVS010000092.1 in S1 Table) respectively. In order to verify if these nonsense mutants lead to similar effects of the deletion mutant, each of these two mutations was introduced into the genome of *P. aeruginosa* PA14. Similar increasement of *cpxP* and *mexA* expression levels and reduction of the antibiotic susceptibility were observed in the PA14*::cpxS^Q85*^*and PA14*::cpxS^E281*^* strains compared to the parental PA14 wild type strain grown in the glucose minimal medium (Fig. 2, Table 1). As these two nonsense mutations were both caused by single nucleotide mutations (^253^CAG→TAG and ^841^GAG→TAG respectively), a lot of single nucleotide mutations of *cpxS* were predictable, at least before the 840^th^ base, to cause nonsense mutations leading to the above cross-talk effects of the development of antibiotic resistance in *P. aeruginosa* PA14 genome (S2 Fig.). Furthermore, it could not be ruled out that some mutants of *cpxS*, other than nonsense mutants, might exist as null mutants. Taken together, *cpxS* null mutations having the risk of multiple way to antibiotic resistance development could be emerged in *P. aeruginosa* clinical isolates.

## Discussion

The regulatory link of CpxR-CpxS two-component system to the expression of the predominant multidrug efflux pump MexAB-OprM underpins the existence of *cpxS* mutational resistome (constitutively active mutants) in *P. aeruginosa*(13). This work has further highlighted that this regulatory link exacerbates antibiotic resistance in multiple way in *P. aeruginosa*. At least 5 EnvZ family sensor kinases can exert cross-talk activation upon CpxR and among them GtrS can lead to increased expression of the MexAB-OprM efflux pump and subsequent reduction of antibiotic susceptibility in *cpxS* null mutants of *P. aeruginosa* under glucose-sole carbon condition respectively (Fig. 2-4, Table 1). Ectopic expression of PmrB is somehow cytotoxic in PA14Δ*cpxS*, so we did not know if PmrB can exert cross-talk activation upon CpxR. It also cannot be ruled out that other EnvZ family sensor kinases stimulated by appropriate physiological signals are able to exert strong enough cross-talk activation upon CpxR leading to increased expression of the MexAB-OprM efflux pump and reduction of antibiotic susceptibility in *cpxS* null mutants of *P. aeruginosa*.

It has been noticed that outnumbered *cpxS* variants versus *cpxR* variants exist among the genomes of *P. aeruginosa* strains including two clinical isolates with nonsense *cpxS* mutants (Fig. 5). Two functionally opposite groups of CpxS variants can exert increased expression of the MexAB-OprM efflux pump and reduction of antibiotic susceptibility: one is constitutively active group reported previously(13) and the other is null mutant group reported in this work. The constitutively active mutants might have fitness cost and be outcompeted in drug-free environment as reported elsewhere(34). In the case of *cpxS* null mutant, the elevated expression of the MexAB-OprM efflux pump and subsequent reduction of antibiotic susceptibility is “hidden”, given that certain signals should trigger the cross-talk effects, probably rendering less fitness cost in *P. aeruginosa*. Since *cpxS* null mutants turn out to be “hidden” yet “multi-bootable” resistome in *P. aeruginosa*, it is appropriate to pay close attention to their impacts in clinical setting.

Contrary to *cpxS* null mutant, *cpxR* null mutant largely reduces the development of antibiotic resistance in *P. aeruginosa*. CpxR strongly accelerates the development of antibiotic resistance during the treatments of sub-lethal concentration antibiotics (Fig. 1). Besides CpxR served as a transcriptional activator, multiple repressors (MexR, NalC and NalD), of which mediating signals need to be fully elucidated, are engaged in the regulation of the expression of the MexAB-OprM efflux pump in *P. aeruginosa*(13). This complicated genetic architecture plus the cross-talk effects described in this work probably render stochastic heterogeneous expression of the MexAB-OprM efflux pump among the population of *P. aeruginosa* cells(35, 36). In this case, CpxR may contribute to hetero-resistance of antibiotics in *P. aeruginosa*. Taking everything into account, CpxR, but not CpxS, should be an important target for combating the development of antibiotic resistance in *P. aeruginosa*.

PmrA-PmrB, another OmpR-EnvZ family two-component system, is related to resistance against cationic antimicrobial peptides, so-called a last-line antibiotic, in *P. aeruginosa*, *K. pneumoniae* and other bacteria(37–41). Numerous *pmrB* variants including nonsense mutants as well as frame-shifted mutants exist in *P. aeruginosa* genomes, outnumbering *pmrA* variants (S1 Fig. F, S1 Table). Accumulating evidences support that PmrB variants are associated with cationic antimicrobial peptides resistance/hetero-resistance(38, 40, 42–44). It is worth to investigate whether either constitutively active PmrB mutants and (or) PmrB null mutants associated with cross-talk effects of other EnvZ sensor kinases lead to cationic antimicrobial peptides resistance/hetero-resistance. Besides PmrA-PmrB, four additional OmpR-EnvZ family two-component systems, PhoP-PhoQ, ParR-ParS, ColR-ColS and CprR-CprS, are involved in cationic antimicrobial peptides resistance in *P. aeruginosa*(45–50). Among them, PhoQ variants are also associated with cationic antimicrobial peptides resistance(45, 50). Again, *phoQ* variants outnumber *phoP* variants in *P. aeruginosa* genomes (S1 Fig. G, S1 Table). It would be rational to investigate cross-talk effects among the relevant OmpR-EnvZ family two-component systems and their impacts on development of cationic antimicrobial peptides resistance in bacteria(51, 52).

Our work emphasizes that the regulatory link of CpxR-CpxS two-component system to the expression of the MexAB-OprM efflux pump is threatening in terms of exacerbating the development of antibiotic resistance. It is worth to note that AdeR-AdeS, a member of OmpR-EnvZ family two-component systems, is divergently located at the upstream of the *adeABC* multidrug efflux pump operon and directly activates its expression in *A. baumannii*(53), another troublesome pathogen prone to develop multidrug resistance(54, 55). Accumulating evidences support that sensor kinase AdeS variants are associated with antibiotic resistance in *A. baumannii*(56–59) and even truncated versions of AdeS caused by IS*Aba1* insertion can activate the expression of the AdeABC multidrug efflux pump through AdeR(60). Probably, more cautious investigations are required, given that multiple mechanisms including cross-talk by other EnvZ sensor kinases may underpin these phenotypes.

In conclusion, our work has provided a new insight to investigate the development of antibiotic resistance exacerbated by OmpR-EnvZ two-component systems and discriminated proper targets for combating antibiotic resistance in *P. aeruginosa* and other bacteria.

## Materials and Methods

### Deletion of gene loci in *P. aeruginosa* strains

Generation of gene-locus-deleted *P. aeruginosa* strains was conducted using a method described previously(10). Briefly, the gene locus of *P. aeruginosa* strains was replaced with a fragment containing the *FRT* gentamicin-resistance (Gm) cassette by double-crossover homologous recombination. The Gm-resistance marker in the chromosome was removed by introducing plasmid harboring the Flp recombinase gene. Correct deletion in the constructed mutant was verified by PCR using primers that bound to flanking chromosomal regions. All DNA primers used in this study are listed in Supplementary Table 2.

### Antibiotic susceptibility test

The MIC of each antibiotic was determined by the two-fold dilution method as previously described(10). Ciprofloxacin and ofloxacin were purchased from Bio Basic Inc. Carbenicillin was purchased from Sigma-Aldrich. Cefsulodin was purchased from TOKU-E (Japan).

### 3-round subcultures exposed to sub-lethal concentration of antibiotics

*P. aeruginosa* cells were grown overnight in LB medium at 37 °C with agitation speed of 180 rpm. They were inoculated 1:1000 in 5 mL of fresh LB medium supplemented with sub-lethal concentration of antibiotics (0.3 μg/mL ofloxacin, MIC 0.5 μg/mL; 60 μg/mL carbenicillin, MIC 128 μg/mL; and no drug control) in 50-mL culture flasks and grown 24 hours at 37 °C with agitation speed of 250 rpm. This subculture procedure was repeated for 3 times. Finally, the cells were serially diluted and 2.5 μL samples were spotted on LB agar plates containing lethal concentration of carbenicillin (200 μg/mL) and ofloxacin (1 μg/mL) to evaluate the development of multidrug resistance.

### β-galactosidase assay for promoter*::lacZ* reporter gene fusion

The *mexA*p::*lacZ* and *cpxP*p*::lacZ* reporter gene fusion plasmids were introduced into *P. aeruginosa* strains by conjugal transfer(10). Cells were grown overnight in MOPS buffered minimal medium(61) supplemented with appropriate nutrients and antibiotics, after which they were inoculated 1:50 in 5 mL of fresh medium in 50-mL culture flasks at 37 °C with agitation speed of 180 rpm. Cells were recovered during the logarithmic growth phase (OD_600_ = 0.5–0.9). β-galactosidase assays were performed as described previously(10). The results are expressed as the mean values from triplicate samples.

### Expression and purification of GtrSc

The plasmid used to express the N-terminal His_6_-tagged GtrSc proteins was constructed by PCR-amplifying the cytoplasmic part of GtrS sequence and cloning it into pET28a (Novagen). The plasmid was transformed into *E. coli* expression host strain BL21(DE3) and grown to an OD_600_ of 0.8 at 37 °C with vigorous shaking in 1 L of LB medium containing kanamycin (50 µg/mL). The cells were then induced with 1 mM IPTG and allowed to express overnight at 22 °C, after which they were harvested by centrifugation. The resulting pellet was resuspended in 10 mL of pre-cooled buffer A (20 mM Tris-HCl, 200 mM NaCl, 1 mM imidazole, pH 8.0) and centrifuged at 4 °C for 10 minutes at 3,500 rpm, after which the pellet was resuspended in 60 mL of pre-cooled buffer A. The cells were disrupted by sonication at 180 W for 8 minutes. The debris and membranes were removed by centrifugation at 4 °C for 60 minutes at 15,000 rpm. The soluble fraction was passed through a 0.2-μm filter and loaded onto a 5-mL nickel column which was previously washed with 10 column volumes of ddH_2_O and equilibrated with 10 column volumes of buffer A. GtrSc proteins were eluted with a mixture of buffer A and buffer B (20 mM Tris-HCl, 200 mM NaCl, 500 mM imidazole, pH 8.0), in which the proportion of buffer B was gradually increased from 0% to 100%. The flow speed was 1 mL/min during the elution process. The protein was collected when its protein peak appeared. The GtrSc protein solution was desalted and concentrated to a final volume of 1.5 mL in buffer C (20 mM Tris-HCl, 200 mM NaCl, pH 8.0). The concentration of GtrSc protein was determined by the Bradford method. GtrSc protein was stored in buffer C supplemented with 50% glycerol at −80 °C.

### Electrophoretic Mobility Shift Assay

The electrophoretic mobility shift assay was carried out using the 2^nd^ generation DIG Gel Shift Kit (Roche)(10). Briefly, 165 bp DNA fragments of the promoter region of *mexA*p was PCR-amplified, after which 150 nM of purified PCR product was DIG-labelled according to the manufacturer’s instructions. 20 μg purified CpxR(10) was phosphorylated with 5 mM ATP by 20 μg purified GtrSc in 100 μl of phosphorylation buffer (50 mM Tris-HCl, 50 mM KCl, 5% glycerol, 5 mM MgCl_2_, 5 mM CaCl_2_, pH 8.0) at room temperature for 1 h. The binding reaction was carried out with 100 nM purified CpxR (with or without GtrSc phosphorylation) and 0.2 nM DIG-labelled DNA fragments at 37 °C for 30 min. The samples were separated by electrophoresis on 6% native polyacrylamide gels, transferred to Hybond-N blotting membranes (Amersham), and visualized by chemiluminescence.

### Construction of *in trans* sensor kinase expression plasmids

In order to control *in trans* sensor kinase expression, a *lacI*^q^-*tac*P region was cloned into broad-host plasmid pBBR1MCS5 to replace the original constitutively expressed *lac* promoter(62). The whole sequence of plasmid vector (pBBR-vector) could be found in supplementary materials. Next, the *in trans* sensor kinase expression systems were constructed by inserting the coding region of each sensor kinase gene into *Nde*I-*Spe*I site using Gibson Assembly method. IPTG (0.2 mM) was added to induce sensor kinase overexpression. It was noted that the basal level of *cpxS* or *gtrS* expression in the absence of IPTG was sufficient to complement the deletion mutants in *P. aeruginosa*.

### Nucleotide substitution in the chromosomal *cpxS*

The target mutant *cpxS* gene fragment was obtained by nest-PCR and introduced into the PA14Δ*cpxS::Gm^R^* strain(13). Following double-crossover homologous recombination, strains sensitive to gentamicin were selected out and verified for successful introduction of the target mutant *cpxS* by sequencing.

## Acknowledgements

This research was supported by the National Science Foundation of China (grant No. 31870129 to Zhe-Xian Tian), the Biological Breeding-National Science and Technology Major Project (grant No. 2024ZD04079 to Zhe-Xian Tian), the National Key R&D Program of China (grant No. 2024YFA091820 to Zhe-Xian Tian), National High Level Hospital Clinical Research Funding (Research Achievement Transformation Project of Peking University First Hospital) (grant No. 2024CX21 to Zhe-Xian Tian), the Sichuan Province Science and Technology Program Project (grant No. 2025YFHZ0060 to Zhe-Xian Tian).

## Supporting Information

**S1 Fig.**
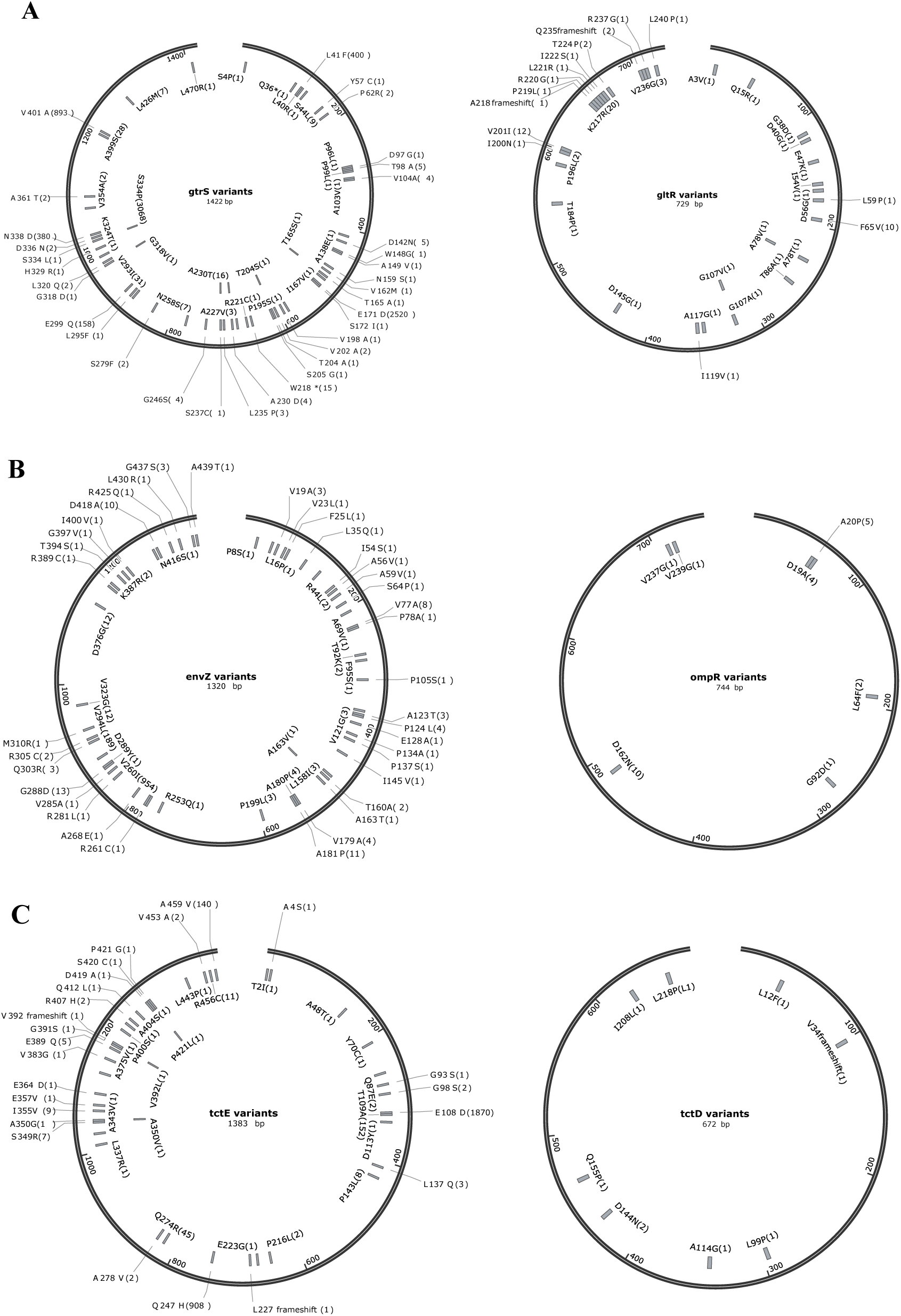
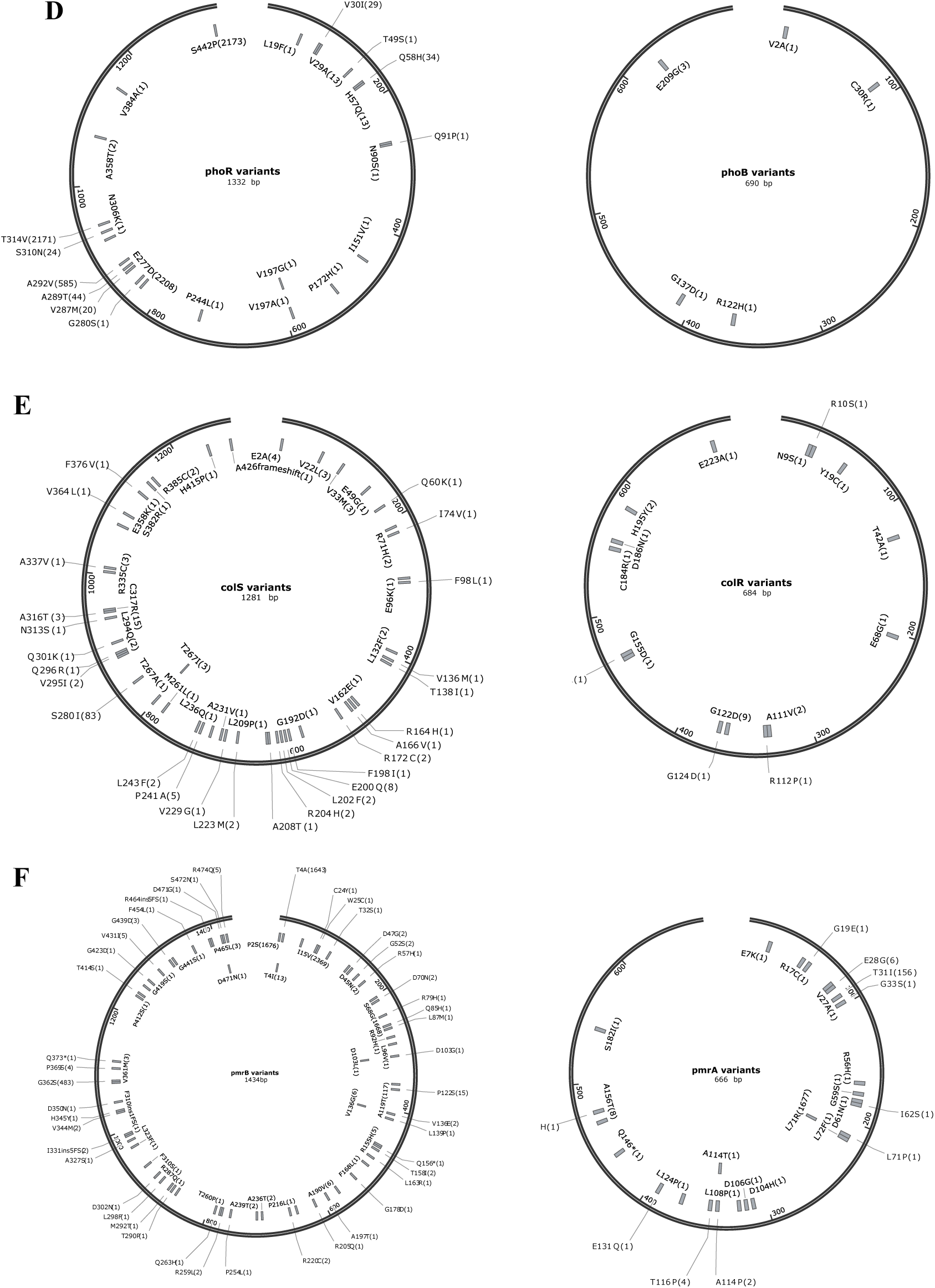
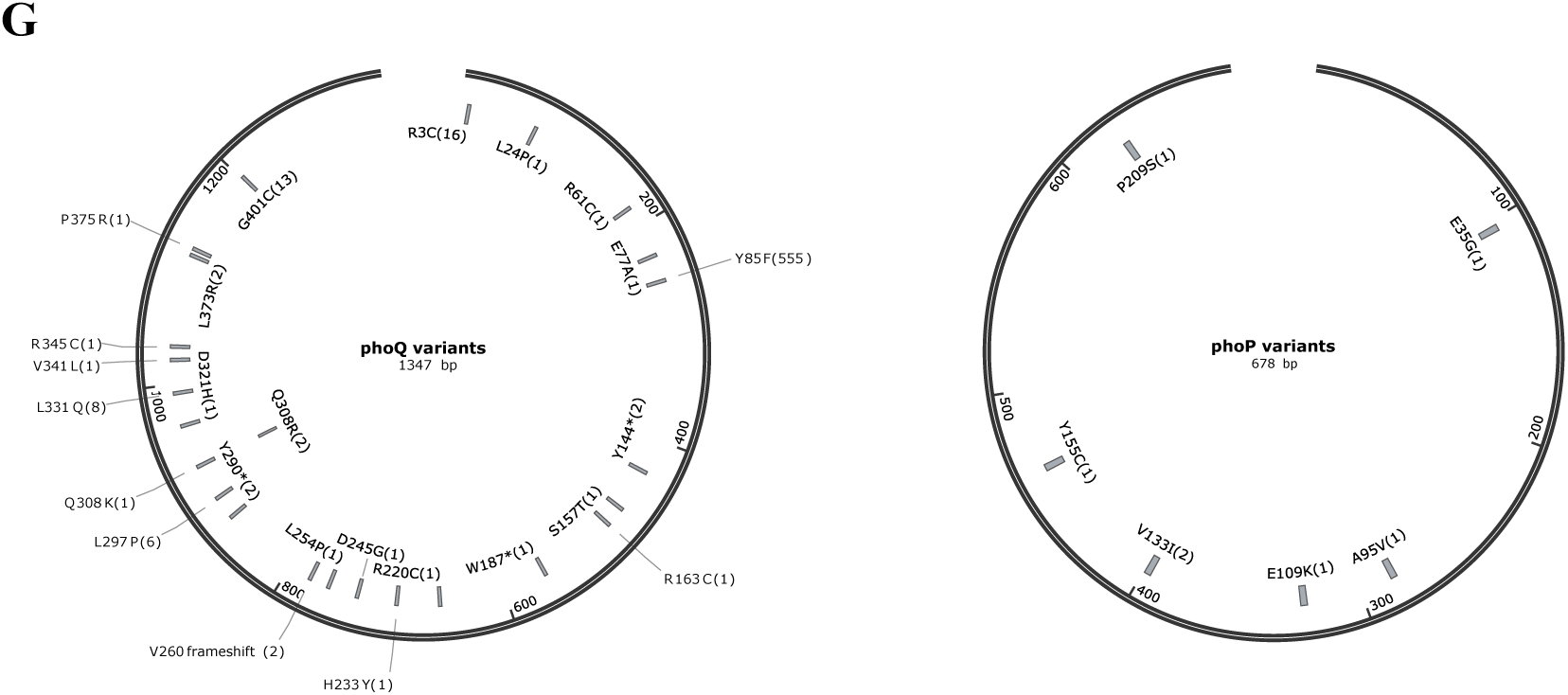
The amino acid substitutions of EnvZ-OmpR family two-components regulatory system variants in the published genomes of *P. aeruginosa* isolates (S1 Table). Numbers in the parentheses are the occurrence times in different genomes. A. GtrS-GltR two-components regulatory system variants; B. EnvZ-OmpR two-components regulatory system variants; C. TctE-TctD two-components regulatory system variants; D. PhoR-PhoB two-components regulatory system variants; E. ColS-ColR two-components regulatory system variants; F. PmrB-PmrA two-components regulatory system variants; G. PhoQ-PhoP two-components regulatory system variants.

**S2 Fig.**
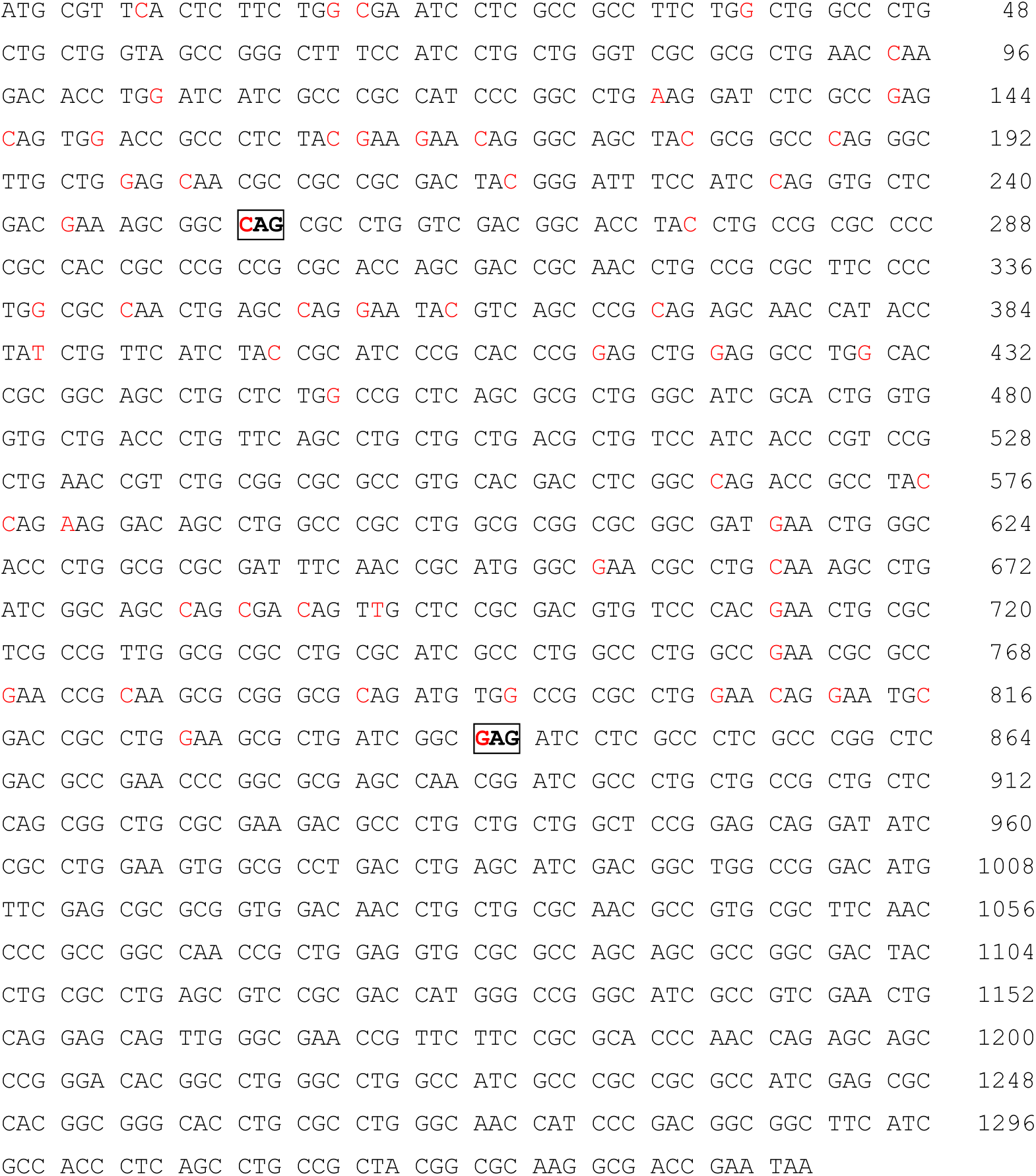
Predicted nucleotide positions of nonsense mutations of the *cpxS* reading frame leading to the cross-talk effects of the development of antibiotic resistance in *P. aeruginosa* PA14. The nucleotides marked in red are able to cause nonsense mutation. The boxed bold codons are the 85^th^ glutamine residue and 281^st^ glutamic acid residue respectively occurred as nonsense mutants in *P. aeruginosa* clinical isolates (Fig. 5).

**S1 Table.** Variants of OmpR family response regulators and EnvZ family sensor kinases among the *P. aeruginosa* genomes.

**S2 Table.** Sequences of DNA primers used in this study.

**S3 Table**. Full sequence of pBBR-vector.

## Notes

### Competing Interest Statement

The authors have declared no competing interest.

